# Individuals with autism have no detectable deficit in neural markers of prediction error when presented with auditory rhythms of varied temporal complexity

**DOI:** 10.1101/2020.05.05.077438

**Authors:** Emily J. Knight, Leona A. Oakes, Susan L. Hyman, Edward G. Freedman, John J. Foxe

## Abstract

The brain’s ability to encode temporal patterns and predict upcoming events is critical for speech perception and other aspects of social communication. Deficits in predictive coding may contribute to difficulties with social communication and overreliance on repetitive predictable environments in individuals with autism spectrum disorder (ASD). Using a mismatch negativity (MMN) task involving rhythmic tone sequences of varying complexity, we tested the hypotheses that 1) individuals with ASD have reduced MMN response to auditory stimuli that deviate in presentation timing from expected patterns, particularly as pattern complexity increases and 2) amplitude of MMN signal is inversely correlated with level of impairment in social communication and repetitive behaviors. Electroencephalography was acquired as individuals (age 6-21years) listened to repeated five-rhythm tones that varied in the Shannon entropy of the rhythm across three conditions (zero, medium-1 bit) and high-2 bit entropy). The majority of the tones conformed to the established rhythm (standard tones); occasionally the 4^th^ tone was temporally shifted relative to its expected time of occurrence (deviant tones). Social communication and repetitive behaviors were measured using the Social Responsiveness Scale and Repetitive Behavior Scale-Revised. Both neurotypical controls (n=19) and individuals with ASD (n=21) show stepwise decreases in MMN as a function of increasing entropy. Contrary to the result forecasted by a predictive coding hypothesis, individuals with ASD do not differ from controls in these neural mechanisms of prediction error to auditory rhythms of varied temporal complexity, and there is no relationship between these signals and social communication or repetitive behavior measures.

**Lay Summary:** We tested the idea that the brain’s ability to use previous experience to influence processing of sounds is weaker in individuals with autism spectrum disorder (ASD) than in neurotypical individuals. We found no difference between individuals with ASD and neurotypical controls in brain wave responses to sounds that occurred earlier than expected in either simple or complex rhythms. There was also no relationship between these brain waves and social communication or repetitive behavior scores.

## INTRODUCTION

Although autism spectrum disorder (ASD) is characterized primarily by deficits in social communication in combination with repetitive behaviors, sensory processing problems are nearly universal in people with ASD (Baum, Stevenson, & Wallace, 2015). Sensory symptoms are among the first symptoms to appear, have been found to be associated with severity of ASD social communication symptoms (Kern et al., 2007), and correlate with attenuated neurophysiological responses during a variety of visual and auditory tasks (Baum et al., 2015; A. B. Brandwein et al., 2015). As a result, they are now a component of the DSM-5 criteria for ASD (American Psychiatric Association, 2013).

In addition to informing our understanding of sensory symptoms and potential development of targeted sensory interventions, understanding differences in the neurophysiologic mechanisms underlying sensory perception may illuminate some aspects of the development of other core features of ASD (Foxe, Molholm, Baudouin, & Wallace, 2018; Jeste & Nelson, 2009). Of particular relevance to the deficits in language-based social interaction manifested by individuals with ASD, multiple aspects of auditory processing appear to be differentially affected in this population (O’Connor, 2012), including orientation and attention to speech and non-speech sounds (Ceponiene et al., 2003; M. A. Dunn, Gomes, & Gravel, 2008; Foxe et al., 2015; Huang et al., 2018; Kuhl, Coffey-Corina, Padden, & Dawson, 2005; Paul, Chawarska, Fowler, Cicchetti, & Volkmar, 2007; Teder-Salejarvi, Pierce, Courchesne, & Hillyard, 2005; Whitehouse & Bishop, 2008), discrimination of sound pitch and duration (Abdeltawwab & Baz, 2015; Chien, Hsieh, & Gau, 2018; Kujala et al., 2007; Lepisto et al., 2005; Lepisto, Nieminen-von Wendt, von Wendt, Naatanen, & Kujala, 2007; Lepisto et al., 2006; Novick, Vaughan, Kurtzberg, & Simson, 1980; Oades, Walker, Geffen, & Stern, 1988; Yu et al., 2015), sensitivity to volume (Khalfa et al., 2004), discrimination of auditory stimuli in the presence of noise (Teder-Salejarvi et al., 2005), processing of speech prosody (Fan & Cheng, 2014; Korpilahti et al., 2007; Kujala et al., 2010; Kujala, Lepisto, Nieminen-von Wendt, Naatanen, & Naatanen, 2005), and response to unexpected or deviant stimuli (Donkers et al., 2015; Ferri et al., 2003; Gomot et al., 2011; Gomot, Giard, Adrien, Barthelemy, & Bruneau, 2002; Hudac et al., 2018; Jansson-Verkasalo et al., 2003; Naatanen, Paavilainen, Rinne, & Alho, 2007; Roberts et al., 2011; Tecchio et al., 2003; Vlaskamp et al., 2017; Weismuller et al., 2015). Neurophysiologic studies of auditory processing in ASD reveal differences in neural activity in response to various speech and non-speech sounds even at very basic levels of sensory processing in the primary auditory cortex; however, these differences may be a result of top-down inputs mediating sensory processing at lower levels (Whitehouse & Bishop, 2008).

Indeed, differences in sensory processing are present not only in auditory but also visual processing, lending credence to the idea that deficits in higher level integrative areas could serve as a unifying explanation for atypical processing across multiple domains (Marco, Hinkley, Hill, & Nagarajan, 2011). Much recent attention has been turned toward a deficit in predictive coding as a primary neurophysiologic mechanism underlying many of the sensory and other symptoms of ASD (Van de Cruys et al., 2014). Predictive coding is the process by which the brain uses previous experience to predict and influence the processing of incoming sensory input via feedback from top down signals to lower level sensory regions (Walsh, McGovern, Clark, & O’Connell, 2020). Many have proposed a deficit in predictive coding in individuals with ASD, whereby there is diminished contribution of top down predictive mechanisms to lower level sensory areas in the processing of incoming auditory and visual stimuli (Pellicano & Burr, 2012; Van de Cruys et al., 2014). Several recent studies in the sensory processing literature have implicated such deficits as a possible explanatory mechanism, and additional commentaries have re-interpreted previous experimental findings in light of this new hypothesis (Baldeweg, 2007;

Garrido, Kilner, Stephan, & Friston, 2009; Wacongne, Changeux, & Dehaene, 2012). Importantly, one of the reasons the predictive coding hypothesis has attracted so much attention is that it may link the extensive basic scientific literature describing deficits in sensory processing of stimuli across multiple domains with the core social communication and repetitive behavior symptoms of ASD. Social communication is highly dependent on efficient multisensory processing and requires extensive rapid application of predictive mechanisms derived from previous experience. Additionally, a relative deficit in predictive coding leads to increased uncertainty in interpreting environmental stimuli and can potentially generate an overreliance on predictability and repetition in the environment (Van de Cruys et al., 2014). As a result, this predictive coding hypothesis of ASD holds great intuitive appeal and theoretical potential. However, there remains limited objective experimental evidence and direct investigation of this hypothesis.

Sensory processing affords an objective and easily manipulated domain within which the predictive coding hypothesis can be rigorously evaluated. Specifically, predictive coding during sensory processing has been invoked as an explanation for the well-described mismatch negativity evoked potential. Mismatch negativity or oddball paradigms involve continuous encephalographic monitoring during presentation of a repeated stimulus interspersed with infrequent oddball or deviant stimuli that differ from the standard stimulus in a key feature (e.g., pitch, duration, location, timing). The neural response to these types of prediction violations manifests as a MMN signal which is a negative deflection in the baseline electroenchalography (EEG) signal typically occurring 150ms-200ms from the onset of the deviant stimulus (Naatanen et al., 2007). The mismatch negativity signal is thought to be related to predictive coding in that it reflects the brain’s ability to generate predictions and detect deviations from these predictions, even during unattended tasks in which the participant is not consciously aware that the stimulus is deviant (Molholm, Martinez, Ritter, Javitt, & Foxe, 2005; Naatanen et al., 2007; Ritter, De Sanctis, Molholm, Javitt, & Foxe, 2006). Furthermore, in typical adults, amplitude of MMN is related to the level of reliability of the auditory stimuli, such that when stimuli are presented in a highly regular manner, a deviant stimulus will generate a stronger MMN than when stimuli are presented in a design with lower predictability (Lumaca, Trusbak Haumann, Brattico, Grube, & Vuust, 2018; Quiroga-Martinez et al., 2019).

There is a large body of literature on other auditory mismatch negativity paradigms in individuals with autism that reports conflicting results. Two recent systematic review and meta-analyses of auditory mismatch negativity studies identified an overall trend toward less pronounced MMN responses to various types of deviant auditory stimuli in individuals with ASD as compared to controls (Chen, Hsieh, Lin, Chan, & Cheng, 2020; Schwartz, Shinn-Cunningham, & Tager-Flusberg, 2018). Yet, these same reviews also described significant variability across studies in the strength and direction of this effect (Chen et al., 2020; Schwartz et al., 2018). An advantage of MMN paradigms is that no explicit behavioral response by the participant to the stimuli is required, thus allowing for the inclusion of individuals with a range of ages and cognitive and verbal abilities. As yet though, the relationship between deficits in MMN responses and core clinical symptoms of ASD such as social communication and repetitive behaviors remains poorly characterized. Additionally, although this theoretical link between mismatch negativity and predictive coding has been proposed (Garrido et al., 2009; Wacongne et al., 2012), no study has specifically tested neural mechanisms of prediction error in individuals with ASD when presented auditory rhythms of varied temporal complexity. If the hypothesis that predictive coding deficits underlie many of the core features of ASD is true, this would make two testable experimental predictions; first, that individuals with ASD should exhibit smaller neurophysiologic responses to prediction error particularly when presented stimuli of higher baseline entropy and, second, that a larger degree of reduction in these responses should correlate with increased symptom severity in the core domains of social communication and restricted interests/repetitive behaviors.

Therefore, the present study aimed to systematically explore the relationship between MMN and the predictive coding of pattern complexity in ASD and to test for association between MMN signal strength and measures of social communication and repetitive behavior in individuals with a range of cognitive and verbal ability. We specifically tested the hypotheses that 1) individuals with ASD have reduced neural MMN responses to auditory stimuli that deviate in presentation timing from expected patterns, particularly as pattern complexity increases and 2) amplitude of MMN signal is inversely correlated with level of impairment in the domains of social communication and restrictive interests/repetitive behaviors in individuals with ASD.

## METHODS

### Participants and Phenotypic Characterization

23 participants with ASD and 21 controls group-matched on age (age range 6-21 years) took part in the study. Participants with ASD were recruited through the University of Rochester Medical Center Developmental and Behavioral Pediatrics Autism Research Registry while typically developing (TD) participants were recruited from the local community. Participants all had normal hearing and normal or corrected-to-normal vision. Participants were excluded if they had a history of traumatic brain injury, history of schizophrenia, bipolar disorder or psychosis, history of neurologic disorder other than well-controlled seizures (as defined by minimal daytime seizure activity with stable medication dose for at least four weeks), or syndromic cause for ASD (e.g. Down’s Syndrome, Fragile X, Tuberous Sclerosis). Participants with history of seizure disorders were included in order to allow for inclusion of more minimally verbal individuals. Participants on psychotropic medications were included. In the ASD group, eleven participants were taking psychotropic medications (selective serotonin or serotonin-norepinephrine reuptake inhibitors (n=4), stimulants (n=3), alpha agonists (n=3), atypical antipsychotics (n=3), atomoxetine (n=2), antiepileptics (n=1), benzodiazepines (n=1), two or more psychotropic medications (n=4). In the control group one participant was taking stimulant medication. All experimental procedures were approved by the institutional review board of the University of Rochester Medical Center. Written informed consent was obtained from the participant or parent/legal guardian. Assent appropriate for age and developmental level was obtained from the participant. Participants were modestly compensated at a rate of $15 per hour for their time in the laboratory.

Individuals with ASD were administered the Autism Diagnostic Observation Schedule-Second Edition (ADOS-2) (Gotham, Risi, Pickles, & Lord, 2007) to validate diagnosis. All participants had a Wechsler Abbreviated Scale of Intelligence (WASI-II) (Wechsler, 2011), and all individuals demonstrated sufficient understanding of the task instructions for this assessment. Receptive language was assessed using the Peabody Picture Vocabulary Test (PPVT-4) (L. Dunn, Dunn, D.M., 2007). Parent questionnaires including Social Responsiveness Scale (SRS-2), (Constantino, 2012) and Repetitive Behavior Scale-Revised (RBS-R) (Lam & Aman, 2007) were administered to all participants to further characterize behavior. Children with anxiety and those with ADHD may also have differences in their responses to unpredicted stimuli. (Cornwell, Garrido, Overstreet, Pine, & Grillon, 2017; Gonzalez-Gadea et al., 2015). Given the high prevalence of these conditions in children with ASD (Avni, Ben-Itzchak, & Zachor, 2018), the Screen for Child Anxiety Related Disorders (SCARED-parent version) (individuals 8-18) (Birmaher et al., 1999) and Swanson, Nolan, and Pelham (SNAP-IV) (individuals 6-18) (Hall et al., 2019) were used to account for these as possible confounders. Detailed medical history was collected for all participants.

Two participants who were initially recruited into the autism group were excluded from the group comparisons, because despite valid autism diagnosis by a qualified professional based on DSM criteria at a younger age, these individuals no longer met instrument classification for ASD on the ADOS-2 and the SRS-2 at time of study. These two participants were included in the neurophysiologic-phenotype correlations which was performed on all participants regardless of group assignment. One participant recruited into the ASD group was withdrawn from the study due to behavioral distress with the electrode cap and gel procedure. No participants in the control group withdrew from the study. Two participants in the ASD group and four participants in the control group had phenotyping assessment performed remotely via videoconference due to Covid-19 institutional isolation regulations. Because the ADOS-2 is not validated for remote administration, ASD diagnosis for remotely assessed participants was confirmed via parental release to access previous ADOS-2 results from the participant’s medical record.

### Stimuli and Task

The experimental paradigm was adapted from that developed by Lumaca et. al. and described in detail in their 2018 publication (Lumaca et al., 2018). Briefly, standard rhythms of five tones were presented repeatedly (70%), interspersed with deviants (30%), in which the fourth tone was shifted 300ms early relative to its standard position. Tones were each sinusoidal with a duration of 50ms, a rise and fall of 5ms, and an intensity of 70(±2)dB SPL. Each five-tone rhythm consisted of tones of a single frequency; however, frequency was randomly varied among three frequencies (315, 397 and 500) across five tone rhythms to prevent habituation. The experiment was divided into three conditions that varied in terms of the complexity of the five-tone rhythm and included a zero-entropy condition in which the tones were evenly spaced, and conditions with medium (1 bit) and high (2 bit) entropy. Entropy was calculated by Shannon entropy: H(X)=-∑*P(xi)*log_2_*P(xi)*) where *x* is the vector of the tone inter-offset-intervals (IOI) in a sequence and *p*(*x_i_*) is the probability that the element *i* occurs in that sequence (given the alphabet of the elements and their relative frequency). Thus, the Shannon entropy is 0 bit for zero-entropy stimuli (400ms IOIs), 1 bit for medium-entropy stimuli (IOIs: 410, 240, 410 and 240ms) and 2 bits for high-entropy stimuli (IOIs: 488, 163, 406 and 244ms). Stimuli were presented with an 800ms interstimulus interval separating the end of each five-tone rhythm from the onset of the next.

The order of standard and deviant stimuli was semi-randomized, with the only constraint that at least one standard pattern appeared before every deviant pattern. Each condition (zero entropy, medium entropy and high entropy) was presented in 2 separate blocks, resulting in six total experimental blocks, each consisting of 400 repeated five-tone rhythms. Block duration was ~16-18 minutes with breaks between as needed between blocks, resulting in ~2hr recording time. The order of conditions was randomized and balanced across subjects. The conditions were counterbalanced within participants such that participants were presented with 1 block each of the zero entropy, medium entropy, and high entropy conditions in random order, followed by presentation of a second block of each condition in reverse order (e.g. medium entropy, zero entropy, high entropy, high entropy, zero entropy, medium entropy).

The task required no specific subject participation, and during the task attention was distracted from the tones by a quiet seated activity of the participant’s choice (e.g. silent movie with subtitles, silent video game, reading a book). An unattended task was chosen to focus on passive generation of neural predictive models, and visual task demands have been shown not to impact the MMN response (Wiens, Szychowska, & Nilsson, 2016). EEG was continuously recorded throughout the task using a 128 channel ActiveTwo acquisition system (BioSemi, The Netherlands). To accommodate a range of developmental needs, we incorporated behavioral strategies including visual schedules, social stories and breaks.

### EEG Acquisition and Preprocessing

All participants sat in a sound-attenuated and electrically-shielded booth (Industrial Acoustics Company, The Bronx, NY). The auditory stimuli were presented to both ears simultaneously via Sennheiser HD 600 open back professional headphones (Sennheiser electronic GmbH & Co. KG, USA). Stimuli were delivered using Presentation^®^ software (Version 18.0, Neurobehavioral Systems, Inc., Berkeley, CA, www.neurobs.com). A Biosemi ActiveTwo (Bio Semi B.V., Amsterdam, Netherlands) 128-electrode array was used to record continuous EEG signals. The set up includes an analog-to-digital converter, and fiber-optic pass-through to a dedicated acquisition computer (digitized at 512 Hz; DC- to-150 Hz pass-band). EEG data were referenced to an active common mode sense electrode and a passive driven right leg electrode.

EEG data were processed and analyzed offline using custom scripts that included functions from the EEGLAB (Delorme & Makeig, 2004) and ERPLAB Toolboxes (Lopez-Calderon & Luck, 2014) for MATLAB (the MathWorks, Natick, MA, USA). EEG data were initially filtered using an IIR Butterworth filter (roll-off 12db/oct, 40db/dec, order 2) with a bandpass set at 0.1–45Hz. Bad channels were manually rejected and interpolated using EEGLAB spherical interpolation. Data were then divided into epochs that started 100ms before the presentation of each tone and extended to 400ms post-stimulus onset. All epochs were then baseline corrected to the 100ms pre-stimulus interval. Trials containing severe movement artifacts or particularly noisy events were rejected if voltages exceeded ±120 μV. The number of interpolated channels and accepted trials for each condition and group is presented in Table S1. The epochs were next averaged as a function of stimulus condition to yield auditory event related potentials (ERP) to the standard and deviant tones. To maximize the relevant ERP signal at fronto-central scalp sites, the data were referenced to a scalp electrode directly posterior and slightly superior to the left mastoid; equivalent to TP9 in the 10-20 system convention). This approach takes advantage of the inversion of the MMN that is seen between frontocentral and mastoid sites (Macdonald & Campbell, 2013).

The window for measurement of the MMN was calculated by subtracting the grand mean ERP to deviant tones from the grand mean ERP to standard tones. The resulting distribution of activity showed maximal difference at ~145ms. We then defined a time window of 10ms centered around 145ms (i.e. 140–150ms) to obtain mean MMN amplitudes for each individual in each condition. Statistical analyses were performed in JASP (JASP Team (2019), Version 0.11.1) with data visualization using interactive dotplot (Weissgerber et al., 2017).

### Data Analyses

The primary analysis employed a repeated-measures analysis of variance (ANOVA) with level of entropy (zero, medium, high) as a within-participant factor and group (ASD vs. TD) as a between-participants factor to examine main effects and their interaction on MMN amplitude. Partial η2 was used to estimate effect sizes. Planned post-hoc tests were used to follow up significant ANOVA effects. A Bayesian repeated measures ANOVA (prior probabilities: r scale fixed effects =0.5; r scale random effects = 1) was also performed in order to compare the marginal likelihoods between the null and alternative hypotheses. A secondary analysis consisting of a repeated-measures ANOVA with SNAP-IV inattention and SCARED anxiety scores as continuous covariates was also performed to assess any confounding impact of these on the results. Data on inattention and ADHD symptoms for individuals outside the validated age range for these phenotyping measures was not included in this secondary analysis.

Stepwise linear regressions assessed the relationship between MMN amplitude in each of the three entropy conditions with scores on the SRS, RBS-R, and PPVT-4.

## RESULTS

Figure 1 displays grand-average ERP waveforms elicited by standard and deviant tones for each level of entropy, as well as the corresponding difference waves, over the fronto-central scalp site (Fz) where the MMN is typically found to show maximal amplitude (Ritter, Sussman, Molholm, & Foxe, 2002). There is an evident stepwise decrease in the amplitude of the MMN response with increasing level of entropy in both the control and ASD groups. These MMNs have typical topography with negativity over the fronto-central sites and positivity over infero-temporal scalp sites bilaterally (Figure 1). MMN lasted from ~50 to 200ms with a maximum at about 145ms, so the mean MMN amplitude within the 140–150ms time-window was chosen for statistical analyses. The mean MMN amplitude for the different entropy conditions in each group is represented in Figure 2. Repeated measures ANOVA revealed a main effect of entropy (F(2,76)=45.906, p=8.465e^-14^, ŋp2=0.547), and post hoc analyses confirmed significant differences between each level of entropy (zero entropy-medium entropy: mean difference= 2.90μV, 95%CI (1.86-3.94), t(39)=7.021,p<4.006e^-8^; zero entropy-high entropy: mean difference= 4.30μV 95%CI (3.02-5.57), t(39)=8.435, p=7.537e^-10^, medium entropy-high entropy: mean difference= 1.39μV, 95%CI (0.32-2.47), t(39)=3.235, p=0.002). However, there was no main effect in the amplitude of the MMN response between the control and ASD group F(1,38)=0.174, p=0.679, η_p_2=0.005), nor was there a group by entropy interaction (F(2,76)=0.239, p=0.788, η_p_2=0.006). Bayesian ANOVA was then performed to assess the marginal likelihood that the data are supportive of the null hypothesis, with a reciprocal Bayes Factor (B01) of 2.140 reflecting that the observed results are about two times more likely to occur under the null hypothesis that there is no difference between individuals with ASD and controls in this measure on a population level. A secondary analysis consisting of a repeated-measures ANOVA with SNAP-IV inattention and SCARED-parent anxiety scores as covariates replicated the presence of a main effect of entropy (F(2,50)=6.601,p=0.004, η_p_2=0.195) and absence of a main effect of group F(1,25)=0.233,p=0.633,η_p_2=0.009), after controlling for comorbid ADHD and anxiety symptoms.

**Figure 1.**
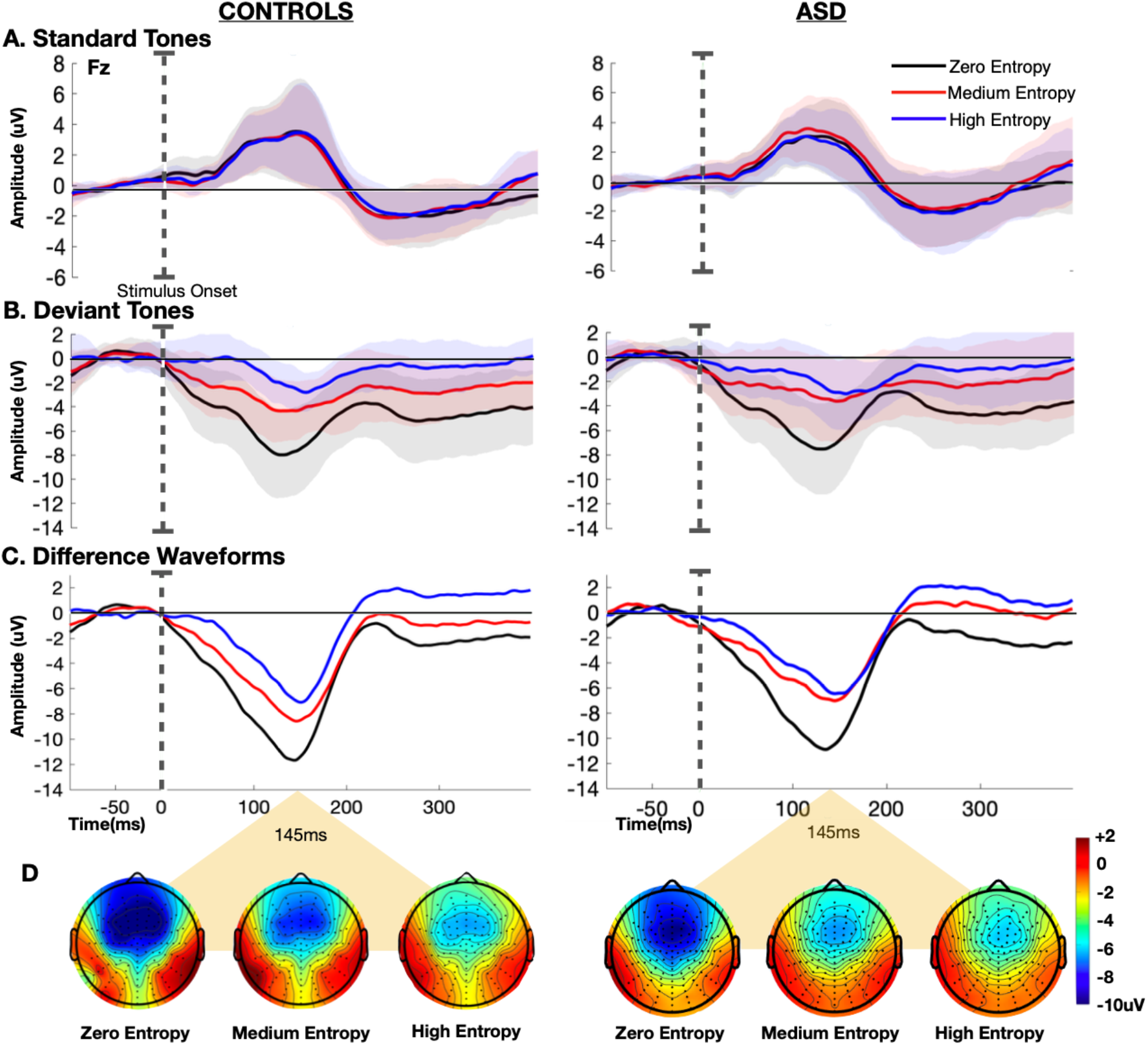
Grand-average event related potentials (ERPs) at electrode Fz obtained in TD controls (n=19) and individuals with ASD (n=21) for **A)** standard tones, **B)** deviant tones, and **C)** difference waveforms (deviant tones-standard tones) in the zero (black), medium (red), and high entropy (blue) conditions. Stimulus onset is marked by the dotted line. Shaded regions mark ± 1 SEM. **D)** Topographic representation of the difference between deviant and standard tones at t=145ms for each of the three conditions in controls (left) and individuals with ASD (right).

**Figure 2.**
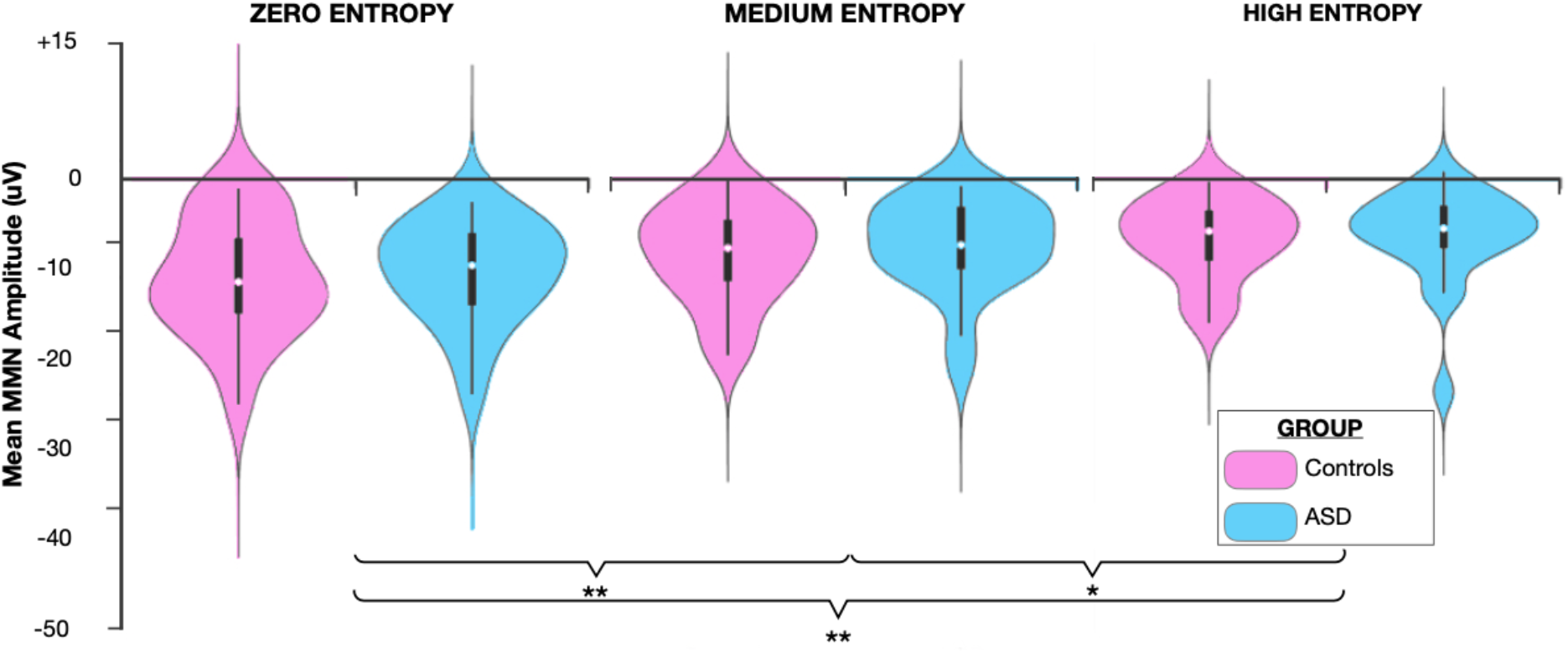
Mean mismatch negativity (MMN) amplitude for each level of entropy in TD controls and individuals with ASD. Significant differences between the levels of entropy are marked by asterisks (*p<0.01, **p<0.001).

Participant characteristics are detailed in Table 1. Mann Whitney U tests were used to assess for group differences on each continuous measure; a chi squared test assessed for group differences in sex. The groups did not differ significantly in age, reflective of appropriate age-matching (U=199.0, p=1.000). The groups did differ in terms of sex distribution with a significantly higher proportion of males in the ASD group (X^2^ (1, n=40) =12.327, p=0.004. No effect of age or sex on MMN was observed in secondary analyses (Figures S1 and S2). Single subject analysis of individuals (n=3) who were minimally verbal as defined by completion of the ADOS2-Module 2 (designed for individuals with phrase speech) suggests intact generation of MMN to this paradigm, although the sample size was insufficient to perform a separate adequately powered group-level analysis on this cohort (Figure S3). Mean IQ in the TD group was higher than that in the ASD group. This difference was related to an effort to be inclusive of individuals with ASD who present with a wide range of cognitive and language abilities. Secondary analysis examining only those children with ASD who have average to above average intelligence and thus better match the control group did not demonstrate a different pattern of findings (Figure S4). As expected, the SRS-2 and RBS-R scores were elevated in the ASD group as compared to controls. Correlation analysis was performed to assess the relationship between the MMN amplitude across the three conditions and SRS-2, RBS-R and PPVT scores, irrespective of group (Figure 3). There was no relationship between MMN amplitude and either SRS-2 (zero entropy: r(40) = 0.027, p=0.432, one-tailed; medium entropy: r(40) = 0.078, p=0.311, one-tailed; high entropy: r(40) = 0.039, p=0.404, one-tailed) or RBS-R scores (zero entropy: r(40) = 0.012, p=0.469, one-tailed; medium entropy: r(40) = 0.027, p=0.433, one-tailed; high entropy: r(40) = 0.085, p=0.29, one-tailed). However, stepwise linear regression revealed a modest relationship between MMN amplitude and receptive language as measured by the PPVT-4, such that higher MMN amplitude in the high entropy condition alone predicted higher receptive language scores (F(1,39) =5.530, p=0.024, R^2^=0.124; (zero entropy: r(40) = −0.205, p=0.100, one-tailed; medium entropy: r(40) = −0.262, p=0.049, one-tailed; high entropy: r(40) = −0.352, p=0.012, one-tailed).

**Figure 3.**
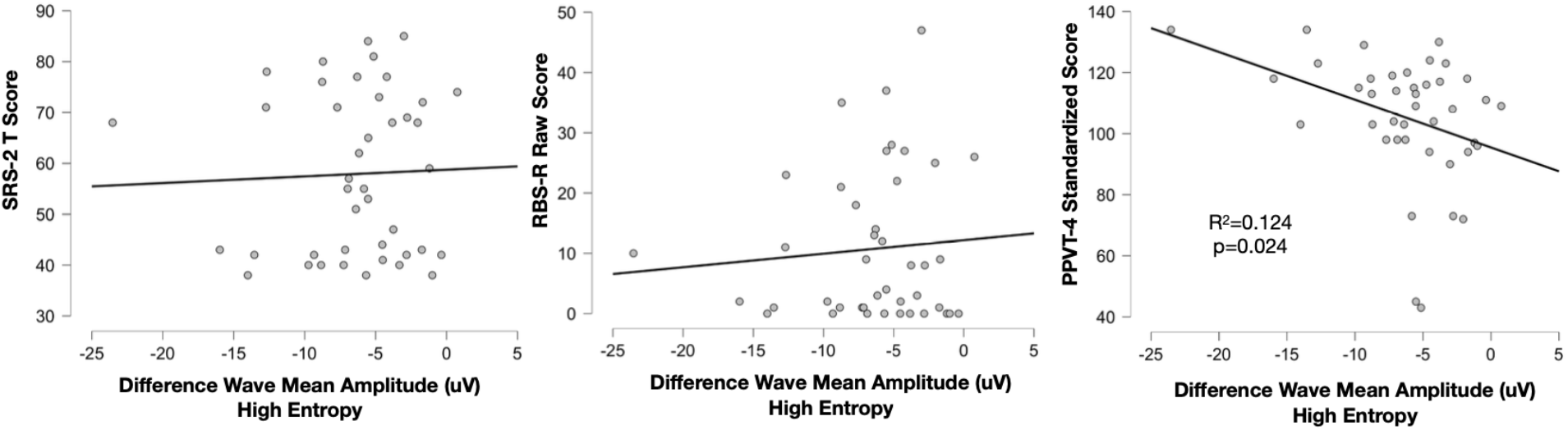
Scatter plots depicting correlation between mean mismatch negativity (MMN) amplitude in the high entropy condition and **A)** SRS-2 T scores **B)** RBS-R raw scores and **C)** PPVT-4 standardized scores. Stepwise linear regression identifies only the high entropy condition as significantly predictive of variance in PPVT-4 scores (R^2^=0.147, p=0.025). Other phenotypic measures show no relationship to MMN amplitude in any of the three entropy conditions.

## DISCUSSION

This study replicates, across a wide developmental age span, findings previously described in neurotypical adults, that the neural response to temporal deviants is inversely proportional to the complexity of the baseline rhythm in which these deviants are embedded (Lumaca et al., 2018). This suggests that the neural mechanisms underlying generation of predictive models of patterns in auditory stimuli and detection of violations to these models are well established by early childhood. Remarkably, these findings were highly replicable across two independent groups of participants in this study irrespective of developmental psychopathology, highlighting the robust nature of these mechanisms.

However, the study tested the specific hypothesis that individuals with ASD would demonstrate a relative weakness in their neural responses to these types of prediction violations as compared to TD controls, consistent with a predictive coding model of ASD. The data obtained provide no support for this model. Furthermore, Bayesian analysis suggests that the findings are at least two times more likely under the null hypothesis that there is no difference between experimental groups on the measures we employed. Thus, it is less likely that the absence of a significant difference is attributable to the study being underpowered; instead this seems to support absence of difference between the two groups in this neurophysiologic mechanism at a population level. Additionally, given the high degree of similarity between groups in the current sample, even if a statistically significant difference emerged with an increased sample size, the absolute magnitude of the effect would be small and have limited clinical utility as a biomarker. Although this does not rule out the involvement of predictive mechanism deficits in other aspects of auditory stimulus processing, multisensory processing or cognition for individuals with ASD, it argues against differences of these mechanisms in the processing of temporal patterns and more broadly against the proposed pervasiveness of predictive coding deficits at the level of basic auditory processing.

MMN responses are fairly widely studied electrophysiological markers in autism neuroscience. Yet, two recent systematic reviews and meta-analyses of these studies highlight significant inconsistency in findings across studies in both the strength and direction of effect, limiting its utility as a biomarker (Chen et al., 2020; Schwartz et al., 2018). In general, both meta-analyses reveal a trend toward decreased MMN responses to auditory stimuli in individuals with ASD, a finding not reflected in the current study. However, none of the studies included in the systematicre views examined MMN responses to timing deviants embedded in rhythms and to our knowledge this is the first study to do so. It is possible that other characteristics of auditory stimuli such as frequency or duration deviants produce more of a differential MMN response in individuals with ASD as compared to TD controls. Other sources of variability across studies include the use of speech vs. pure tone stimuli and heterogeneity in the age or symptom profile of the included subject population. Additionally, sample sizes are often small with sample sizes frequently less than ten per group, increasing the likelihood of skewed samples that are less representative of the population as a whole. Importantly, individuals who are minimally verbal or nonverbal are generally underrepresented in existing studies.

Although there was individual variability in the amplitude of MMN within both groups (ASD and controls), we found no relationship between an individual’s pattern of MMN responses and phenotypic characteristics including social communication and repetitive behaviors. There was a weak positive correlation between MMN amplitude and PPVT-4 score suggesting a possible relationship between neural responses to timing deviants and receptive vocabulary; however, further experimentation including individuals with a wider range of receptive language ability would be necessary to determine the reliability of this finding. This study also addressed a limitation in many existing studies of sensory perception and predictive processes in individuals with ASD, which is the possible influence of comorbid conditions as drivers of differences detected between ASD and control groups. As expected, rates of inattention and anxiety were both higher in the ASD than in the control group in the current study; however, there appeared to be no influence of these potential confounders on the results.

### Study Limitations

This study sample included a fairly heterogenous population, with variability across several demographic and phenotypic dimensions. A potential limitation of the current study is the wide age-range of the participants. Previous studies have suggested that auditory responses continue to mature with typical development across the age-range tested (Bishop, Anderson, Reid, & Fox, 2011; Bishop, Hardiman, & Barry, 2011; Alice B. Brandwein et al., 2011). However, secondary analyses of the data from the present study addressing age as a covariate did not reveal any developmental differences in markers of neural prediction error with age. Consistent with the ASD population sex distribution (Kogan et al., 2018), there was a male predominance in the ASD group that was not reflected in the control group; we did not specifically match groups based on sex due to reports that sex does not influence auditory MMN (Kasai et al., 2002), and secondary analysis comparing males and females within the study control group also do not demonstrate sex differences. There was also a significant difference in IQ between the ASD and control groups due to the inclusion of three minimally verbal individuals who had below average IQ in the ASD group. Again, secondary analyses examining only those children with ASD who have average to above average intelligence and thus better match the control group did not suggest an influential role for these factors. Furthermore, given the lack of a detectable difference in MMN responses between the ASD and control groups, it is unlikely that these group differences in sex or IQ impacted our conclusions.

Finally, the majority of the patient sample consisted of individuals with average to above average intelligence and fluent language. Representation of individuals with limited language was insufficient to perform adequately powered statistical analysis examining this population specifically; however, single subject analysis of these individuals revealed intact MMN generation to deviants within each level of complexity.

### Conclusions

In conclusion, we were able to adapt an experimental paradigm that modulates the level of entropy of an auditory rhythm in order to directly test the contribution of predictive coding mechanisms to processing of auditory stimulus temporal characteristics in individuals with ASD. We found no difference between individuals with ASD and TD controls in neural mechanisms of prediction error when presented with auditory rhythms of varied temporal complexity, providing experimental evidence against a significant role for predictive coding deficits in this type of processing in ASD.

## ACKNOWLEDGEMENTS

The authors acknowledge the contribution of undergraduate students: Dalia Mitchel, Zhewei Cao, Oren Bazer, and Rose Cash and graduate student Kathryn Toffolo, B.S. for their assistance with data collection. This work was supported by the Ernest J. Del Monte Institute for Neuroscience Pilot Program for 2019 via the Harry T. Mangurian Foundation. The work of Dr. Knight was supported through the University of Rochester Medical Center Department of Pediatrics Chair Fellow Award and a Bradford Fellowship.

## AUTHOR CONTRIBUTIONS

JJF, EK and EGF conceived the study and designed the original experiment. EK coordinated data collection, analyzed the data, and constructed the illustrations. SH provided oversight for development of theoretical considerations related to the patient population and phenotyping plan and facilitated participant recruitment. LA took primary responsibility for coordinating recruitment and phenotyping of participants. EK wrote the first draft of the manuscript. All authors provided editorial input to EK on subsequent drafts. All authors read and approved the final version of this manuscript.

## DATA SHARING

Upon acceptance for publication, the authors will coordinate with the editorial office to ensure that the full and appropriately de-identified datasets are uploaded to a publicly available repository and that the appropriate link is made to the article file.

## COMPLIANCE WITH ETHICAL STANDARDS

The Research Subjects Review Board of the University of Rochester approved all the experimental procedures (STUDY00002036). Each participant provided written informed consent in accordance with the tenets laid out in the Declaration of Helsinki.

## Abbreviations

ASD: autism spectrum disorder
TD: typically developing
MMN: mismatch negativity
EEG: electroencephalography

## REFERENCES

Abdeltawwab, M. M., & Baz, H. (2015). Automatic Pre-Attentive Auditory Responses: MMN to Tone Burst Frequency Changes in Autistic School-Age Children. J Int Adv Otol, 11(1), 36–41. doi:10.5152/iao.2014.438

American Psychiatric Association. (2013). Diagnostic and Statistical Manual of Mental Disorders (5th ed.). Washington, DC.

Avni, E., Ben-Itzchak, E., & Zachor, D. A. (2018). The Presence of Comorbid ADHD and Anxiety Symptoms in Autism Spectrum Disorder: Clinical Presentation and Predictors. Front Psychiatry, 9, 717. doi:10.3389/fpsyt.2018.00717

Baldeweg, T. (2007). ERP Repetition Effects and Mismatch Negativity Generation. Journal of Psychophysiology, 21(3-4), 204–213. doi:10.1027/0269-8803.21.34.204

Baum, S. H., Stevenson, R. A., & Wallace, M. T. (2015). Behavioral, perceptual, and neural alterations in sensory and multisensory function in autism spectrum disorder. Prog Neurobiol, 134, 140–160. doi:10.1016/j.pneurobio.2015.09.007

Birmaher, B., Brent, D. A., Chiappetta, L., Bridge, J., Monga, S., & Baugher, M. (1999). Psychometric properties of the Screen for Child Anxiety Related Emotional Disorders (SCARED): a replication study. J Am Acad Child Adolesc Psychiatry, 38(10), 1230–1236. doi:10.1097/00004583-199910000-00011

Bishop, D. V., Anderson, M., Reid, C., & Fox, A. M. (2011). Auditory development between 7 and 11 years: an event-related potential (ERP) study. PLoS One, 6(5), e18993. doi:10.1371/journal.pone.0018993

Bishop, D. V., Hardiman, M. J., & Barry, J. G. (2011). Is auditory discrimination mature by middle childhood? A study using time-frequency analysis of mismatch responses from 7 years to adulthood. Dev Sci, 14(2), 402–416. doi:10.1111/j.1467-7687.2010.00990.x

Brandwein, A. B., Foxe, J. J., Butler, J. S., Frey, H. P., Bates, J. C., Shulman, L. H., & Molholm, S. (2015). Neurophysiological indices of atypical auditory processing and multisensory integration are associated with symptom severity in autism. J Autism Dev Disord, 45(1), 230–244. doi:10.1007/s10803-014-2212-9

Brandwein, A. B., Foxe, J. J., Russo, N. N., Altschuler, T. S., Gomes, H., & Molholm, S. (2011). The development of audiovisual multisensory integration across childhood and early adolescence: A high-density electrical mapping study. Cerebral Cortex, 21(5), 1042–1055. doi:10.1093/cercor/bhq170

Ceponiene, R., Lepisto, T., Shestakova, A., Vanhala, R., Alku, P., Naatanen, R., & Yaguchi, K. (2003). Speech-sound-selective auditory impairment in children with autism: they can perceive but do not attend. Proc Natl Acad Sci USA, 100(9), 5567–5572. doi:10.1073/pnas.0835631100

Chen, T. C., Hsieh, M. H., Lin, Y. T., Chan, P. S., & Cheng, C. H. (2020). Mismatch negativity to different deviant changes in autism spectrum disorders: A meta-analysis. Clin Neurophysiol, 131(3), 766–777. doi:10.1016/j.clinph.2019.10.031

Chien, Y. L., Hsieh, M. H., & Gau, S. S. (2018). Mismatch Negativity and P3a in Adolescents and Young Adults with Autism Spectrum Disorders: Behavioral Correlates and Clinical Implications. J Autism Dev Disord, 48(5), 1684–1697. doi:10.1007/s10803-017-3426-4

Constantino, J. N., Gruber, C. P.. (2012). Social Responsiveness Scale, Second Edition. Los Angeles, CA: Western Psychological Services.

Cornwell, B. R., Garrido, M. I., Overstreet, C., Pine, D. S., & Grillon, C. (2017). The Unpredictive Brain Under Threat: A Neurocomputational Account of Anxious Hypervigilance. Biol Psychiatry, 82(6), 447–454. doi:10.1016/j.biopsych.2017.06.031

Delorme, A., & Makeig, S. (2004). EEGLAB: an open source toolbox for analysis of single-trial EEG dynamics including independent component analysis. J Neurosci Methods, 134(1), 9–21. doi:10.1016/j.jneumeth.2003.10.009

Donkers, F. C., Schipul, S. E., Baranek, G. T., Cleary, K. M., Willoughby, M. T., Evans, A. M., … Belger, A. (2015). Attenuated auditory event-related potentials and associations with atypical sensory response patterns in children with autism. J Autism Dev Disord, 45(2), 506–523. doi:10.1007/s10803-013-1948-y

Dunn, L., Dunn, D. M. (2007). PPVT-4 :Peabody picture vocabulary test. Minneapolis, MN: Pearson Assessments.

Dunn, M. A., Gomes, H., & Gravel, J. (2008). Mismatch negativity in children with autism and typical development. J Autism Dev Disord, 38(1), 52–71. doi:10.1007/s10803-007-0359-3

Fan, Y. T., & Cheng, Y. (2014). Atypical mismatch negativity in response to emotional voices in people with autism spectrum conditions. PLoS One, 9(7), e102471. doi:10.1371/journal.pone.0102471

Ferri, R., Elia, M., Agarwal, N., Lanuzza, B., Musumeci, S. A., & Pennisi, G. (2003). The mismatch negativity and the P3a components of the auditory event-related potentials in autistic low-functioning subjects. Clin Neurophysiol, 114(9), 1671–1680. doi:10.1016/s1388-2457(03)00153-6

Foxe, J. J., Molholm, S., Baudouin, S. J., & Wallace, M. T. (2018). Explorations and perspectives on the neurobiological bases of autism spectrum disorder. Eur JNeurosci, 47(6), 488–496. doi:10.1111/ejn.13902

Foxe, J. J., Molholm, S., Del Bene, V. A., Frey, H. P., Russo, N. N., Blanco, D., … Ross, L. A. (2015). Severe multisensory speech integration deficits in high-functioning school-aged children with autism spectrum disorder (ASD) and their resolution during early adolescence. Cerebral Cortex, 25(2), 298–312. doi:10.1093/cercor/bht213

Garrido, M. I., Kilner, J. M., Stephan, K. E., & Friston, K. J. (2009). The mismatch negativity: a review of underlying mechanisms. Clin Neurophysiol, 120(3), 453–463. doi:10.1016/j.clinph.2008.11.029

Gomot, M., Blanc, R., Clery, H., Roux, S., Barthelemy, C., & Bruneau, N. (2011). Candidate electrophysiological endophenotypes of hyper-reactivity to change in autism. J Autism Dev Disord, 41(6), 705–714. doi:10.1007/s10803-010-1091-y

Gomot, M., Giard, M. H., Adrien, J. L., Barthelemy, C., & Bruneau, N. (2002). Hypersensitivity to acoustic change in children with autism: electrophysiological evidence of left frontal cortex dysfunctioning. Psychophysiology, 39(5), 577–584. doi:10.1017.S0048577202394058

Gonzalez-Gadea, M. L., Chennu, S., Bekinschtein, T. A., Rattazzi, A., Beraudi, A., Tripicchio, P., … Ibanez, A. (2015). Predictive coding in autism spectrum disorder and attention deficit hyperactivity disorder. J Neurophysiol, 114(5), 2625–2636. doi:10.1152/jn.00543.2015

Gotham, K., Risi, S., Pickles, A., & Lord, C. (2007). The Autism Diagnostic Observation Schedule: revised algorithms for improved diagnostic validity. J Autism Dev Disord, 37(4), 613–627. doi:10.1007/s10803-006-0280-1

Hall, C. L., Guo, B., Valentine, A. Z., Groom, M. J., Daley, D., Sayal, K., & Hollis, C. (2019). The Validity of the SNAP-IV in Children Displaying ADHD Symptoms. Assessment, 1073191119842255. doi:10.1177/1073191119842255

Huang, D., Yu, L., Wang, X., Fan, Y., Wang, S., & Zhang, Y. (2018). Distinct patterns of discrimination and orienting for temporal processing of speech and nonspeech in Chinesechildren with autism: an event-related potential study. Eur J Neurosci, 47(6), 662–668. doi:10.1111/ejn.13657

Hudac, C. M., DesChamps, T. D., Arnett, A. B., Cairney, B. E., Ma, R., Webb, S. J., & Bernier, R. A. (2018). Early enhanced processing and delayed habituation to deviance sounds in autism spectrum disorder. Brain Cogn, 123, 110–119. doi:10.1016/j.bandc.2018.03.004

Jansson-Verkasalo, E., Ceponiene, R., Kielinen, M., Suominen, K., Jantti, V., Linna, S. L., … Naatanen, R. (2003). Deficient auditory processing in children with Asperger Syndrome, as indexed by event-related potentials. Neurosci Lett, 338(3), 197–200. doi:10.1016/s0304-3940(02)01405-2

Jeste, S. S., & Nelson, C. A., 3rd. (2009). Event related potentials in the understanding of autism spectrum disorders: an analytical review. J Autism Dev Disord, 39(3), 495–510. doi:10.1007/s10803-008-0652-9

Kasai, K., Nakagome, K., Iwanami, A., Fukuda, M., Itoh, K., Koshida, I., & Kato, N. (2002). No effect of gender on tonal and phonetic mismatch negativity in normal adults assessed by a high-resolution EEG recording. Brain Res Cogn Brain Res, 13(3), 305–312. doi:10.1016/s0926-6410(01)00125-2

Kern, J. K., Trivedi, M. H., Grannemann, B. D., Garver, C. R., Johnson, D. G., Andrews, A. A., … Schroeder, J. L. (2007). Sensory correlations in autism. Autism, 11(2), 123–134. doi:10.1177/1362361307075702

Khalfa, S., Bruneau, N., Roge, B., Georgieff, N., Veuillet, E., Adrien, J. L., … Collet, L. (2004). Increased perception of loudness in autism. Hear Res, 198(1-2), 87–92. Retrieved from https://www.ncbi.nlm.nih.gov/pubmed/15617227

Kogan, M. D., Vladutiu, C. J., Schieve, L. A., Ghandour, R. M., Blumberg, S. J., Zablotsky, B., … Lu, M. C. (2018). The Prevalence of Parent-Reported Autism Spectrum Disorder Among US Children. Pediatrics, 142(6). doi:10.1542/peds.2017-4161

Korpilahti, P., Jansson-Verkasalo, E., Mattila, M. L., Kuusikko, S., Suominen, K., Rytky, S., … Moilanen, I. (2007). Processing of affective speech prosody is impaired in Asperger syndrome. J Autism Dev Disord, 37(8), 1539–1549. doi:10.1007/s10803-006-0271-2

Kuhl, P. K., Coffey-Corina, S., Padden, D., & Dawson, G. (2005). Links between social and linguistic processing of speech in preschool children with autism: behavioral and electrophysiological measures. Dev Sci, 8(1), F1–F12. doi:10.1111/j.1467-7687.2004.00384.x

Kujala, T., Aho, E., Lepisto, T., Jansson-Verkasalo, E., Nieminen-von Wendt, T., von Wendt, L., & Naatanen, R. (2007). Atypical pattern of discriminating sound features in adults with Asperger syndrome as reflected by the mismatch negativity. Biol Psychol, 75(1), 109–114. doi:10.1016/j.biopsycho.2006.12.007

Kujala, T., Kuuluvainen, S., Saalasti, S., Jansson-Verkasalo, E., Wendt, L. V., & Lepisto, T. (2010). Speech-feature discrimination in children with Asperger syndrome as determined with the multi-feature mismatch negativity paradigm. Clin Neurophysiol, 121(9), 1410–1419. doi:10.1016/j.clinph.2010.03.017

Kujala, T., Lepisto, T., Nieminen-von Wendt, T., Naatanen, P., & Naatanen, R. (2005). Neurophysiological evidence for cortical discrimination impairment of prosody in Asperger syndrome. Neurosci Lett, 383(3), 260–265. doi:10.1016/j.neulet.2005.04.048

Lam, K. S., & Aman, M. G. (2007). The Repetitive Behavior Scale-Revised: independent validation in individuals with autism spectrum disorders. J Autism Dev Disord, 37(5), 855–866. doi:10.1007/s10803-006-0213-z

Lepisto, T., Kujala, T., Vanhala, R., Alku, P., Huotilainen, M., & Naatanen, R. (2005). The discrimination of and orienting to speech and non-speech sounds in children with autism. Brain Res, 1066(1-2), 147–157. doi:10.1016/j.brainres.2005.10.052

Lepisto, T., Nieminen-von Wendt, T., von Wendt, L., Naatanen, R., & Kujala, T. (2007). Auditory cortical change detection in adults with Asperger syndrome. Neurosci Lett, 414(2), 136-140. doi:10.1016/j.neulet.2006.12.009

Lepisto, T., Silokallio, S., Nieminen-von Wendt, T., Alku, P., Naatanen, R., & Kujala, T. (2006). Auditory perception and attention as reflected by the brain event-related potentials in children with Asperger syndrome. Clin Neurophysiol, 117(10), 2161–2171. doi:10.1016/j.clinph.2006.06.709

Lopez-Calderon, J., & Luck, S. J. (2014). ERPLAB: an open-source toolbox for the analysis of event-related potentials. Front Hum Neurosci, 8, 213. doi:10.3389/fnhum.2014.00213

Lumaca, M., Trusbak Haumann, N., Brattico, E., Grube, M., & Vuust, P. (2018). Weighting of neural prediction error by rhythmic complexity: a predictive coding account using Mismatch Negativity. Eur J Neurosci. doi:10.1111/ejn.14329

Macdonald, M., & Campbell, K. (2013). Event-related potential measures of a violation of an expected increase and decrease in intensity. PLoS One, 8(10), e76897. doi:10.1371/journal.pone.0076897

Marco, E. J., Hinkley, L. B., Hill, S. S., & Nagarajan, S. S. (2011). Sensory processing in autism: a review of neurophysiologic findings. Pediatr Res, 69(5 Pt 2), 48R–54R. doi:10.1203/PDR.0b013e3182130c54

Molholm, S., Martinez, A., Ritter, W., Javitt, D. C., & Foxe, J. J. (2005). The neural circuitry of preattentive auditory change-detection: an fMRI study of pitch and duration mismatch negativity generators. Cereb Cortex, 15(5), 545–551. doi:10.1093/cercor/bhh155

Naatanen, R., Paavilainen, P., Rinne, T., & Alho, K. (2007). The mismatch negativity (MMN) in basic research of central auditory processing: a review. Clin Neurophysiol, 118(12), 2544–2590. doi:10.1016/j.clinph.2007.04.026

Novick, B., Vaughan, H. G., Jr., Kurtzberg, D., & Simson, R. (1980). An electrophysiologic indication of auditory processing defects in autism. Psychiatry Res, 3(1), 107–114. doi:10.1016/0165-1781(80)90052-9

O’Connor, K. (2012). Auditory processing in autism spectrum disorder: a review. Neurosci Biobehav Rev, 36(2), 836–854. doi:10.1016/j.neubiorev.2011.11.008

Oades, R. D., Walker, M. K., Geffen, L. B., & Stern, L. M. (1988). Event-related potentials in autistic and healthy children on an auditory choice reaction time task. Int J Psychophysiol, 6(1), 25–37. doi:10.1016/0167-8760(88)90032-3

Paul, R., Chawarska, K., Fowler, C., Cicchetti, D., & Volkmar, F. (2007). “Listen my children and you shall hear”: auditory preferences in toddlers with autism spectrum disorders. J Speech Lang Hear Res, 50(5), 1350–1364. doi:10.1044/1092-4388(2007/094)

Pellicano, E., & Burr, D. (2012). When the world becomes ‘too real’: a Bayesian explanation of autistic perception. Trends Cogn Sci, 16(10), 504–510. doi:10.1016/j.tics.2012.08.009

Quiroga-Martinez, D. R., Hansen, N. C., Hojlund, A., Pearce, M. T., Brattico, E., & Vuust, P. (2019). Reduced prediction error responses in high-as compared to low-uncertainty musical contexts. Cortex, 120, 181–200. doi:10.1016/j.cortex.2019.06.010

Ritter, W., De Sanctis, P., Molholm, S., Javitt, D. C., & Foxe, J. J. (2006). Preattentively grouped tones do not elicit MMN with respect to each other. Psychophysiology, 43(5), 423–430. doi:10.1111/j.1469-8986.2006.00423.x

Ritter, W., Sussman, E., Molholm, S., & Foxe, J. J. (2002). Memory reactivation or reinstatement and the mismatch negativity. Psychophysiology, 39(2), 158–165. doi:10.1017/S0048577202001622

Roberts, T. P., Cannon, K. M., Tavabi, K., Blaskey, L., Khan, S. Y., Monroe, J. F. Edgar, J. C. (2011). Auditory magnetic mismatch field latency: a biomarker for language impairment in autism. Biol Psychiatry, 70(3), 263–269. doi:10.1016/j.biopsych.2011.01.015

Schwartz, S., Shinn-Cunningham, B., & Tager-Flusberg, H. (2018). Meta-analysis and systematic review of the literature characterizing auditory mismatch negativity in individuals with autism. Neurosci Biobehav Rev, 87, 106–117. doi:10.1016/j.neubiorev.2018.01.008

Tecchio, F., Benassi, F., Zappasodi, F., Gialloreti, L. E., Palermo, M., Seri, S., & Rossini, P. M. (2003). Auditory sensory processing in autism: a magnetoencephalographic study. Biol Psychiatry, 54(6), 647–654. doi:10.1016/s0006-3223(03)00295-6

Teder-Salejarvi, W. A., Pierce, K. L., Courchesne, E., & Hillyard, S. A. (2005). Auditory spatial localization and attention deficits in autistic adults. Brain Res Cogn Brain Res, 23(2-3), 221–234. doi:10.1016/j.cogbrainres.2004.10.021

Van de Cruys, S., Evers, K., Van der Hallen, R., Van Eylen, L., Boets, B., de-Wit, L., & Wagemans, J. (2014). Precise minds in uncertain worlds: predictive coding in autism. Psychol Rev, 121(4), 649–675. doi:10.1037/a0037665

Vlaskamp, C., Oranje, B., Madsen, G. F., Mollegaard Jepsen, J. R., Durston, S., Cantio, C., Bilenberg, N. (2017). Auditory processing in autism spectrum disorder: Mismatch negativity deficits. Autism Res, 1θ(n), 1857–1865. doi:10.1002/aur.1821

Wacongne, C., Changeux, J. P., & Dehaene, S. (2012). A neuronal model of predictive coding accounting for the mismatch negativity. J Neurosci, 32(11), 3665–3678. doi:10.1523/JNEUROSCI.5003-11.2012

Walsh, K. S., McGovern, D. P., Clark, A., & O’Connell, R. G. (2020). Evaluating the neurophysiological evidence for predictive processing as a model of perception. Ann N Y Acad Sci. doi:10.1111/nyas.14321

Wechsler, D. (2011). Wechsler Abbreviated Scale of Intelligence-2nd Edition. San Antonio, TX: Psychological Corporation.

Weismuller, B., Thienel, R., Youlden, A. M., Fulham, R., Koch, M., & Schall, U. (2015). Psychophysiological Correlates of Developmental Changes in Healthy and Autistic Boys. J Autism Dev Disord, 45(7), 2168–2175. doi:10.1007/s10803-015-2385-x

Weissgerber, T. L., Savic, M., Winham, S. J., Stanisavljevic, D., Garovic, V. D., & Milic, N. M. (2017). Data visualization, bar naked: A free tool for creating interactive graphics. J Biol Chem, 292(50), 20592–20598. doi:10.1074/jbc.RA117.000147

Whitehouse, A. J. O., & Bishop, D. V. M. (2008). Do children with autism ‘switch off’ to speech sounds? An investigation using event-related potentials. Developmental Science, 11(4), 516–524. doi:10.1111/j.1467-7687.2008.00697.x

Wiens, S., Szychowska, M., & Nilsson, M. E. (2016). Visual Task Demands and the Auditory Mismatch Negativity: An Empirical Study and a Meta-Analysis. PLoS One, 11(1), e0146567. doi:10.1371/journal.pone.0146567

Yu, L., Fan, Y., Deng, Z., Huang, D., Wang, S., & Zhang, Y. (2015). Pitch Processing in Tonal-Language-Speaking Children with Autism: An Event-Related Potential Study. J Autism Dev Disord, 45(11), 3656–3667. doi:10.1007/s10803-015-2510-x

