## Supplemental Table and Figures for "Individuals with autism have no detectable deficit in neural markers of prediction error when presented with auditory rhythms of varied temporal complexity"

**Table S1.** Numbers of trial included in the analysis across condition in controls and in participants with ASD.

| | TD (n)=19 | | | ASD (n)=21 | | | TD All Accepted | ASD All Accepted |
|---|---|---|---|---|---|---|---|---|
| | **Zero Entropy** | **Medium Entropy** | **High Entropy** | **Zero Entropy** | **Medium Entropy** | **High Entropy** | **Across Conditions** | **Across Conditions** |
| Avg. Accepted Standard trials ± SD | 485±100 | 484±70 | 445±96 | 415±110 | 428±102 | 412±105 | 472±90 | 418±104 |
| Avg. Accepted Deviant trials ± SD | 205±44 | 203±31 | 187±39 | 174±49 | 185±42 | 170±47 | 198±38 | 176±46 |
| Median Channels Interpolated (range) | | | | | | | 1 (0-10) | 0 (0-14) |

**S1. Difference Waveforms by Age**

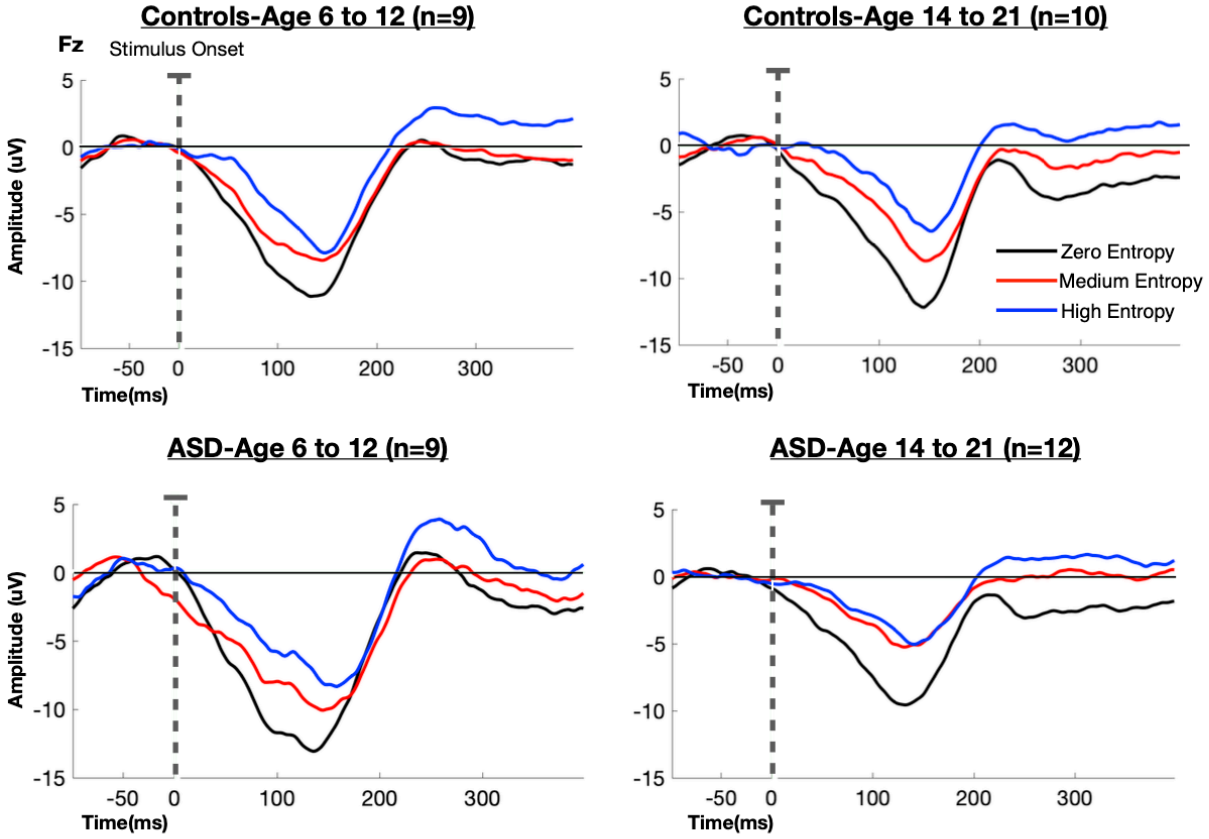


**S1.** Grand-average difference waveforms (deviant tones-standard tones) for age groups 6-12 years and 14-21 years in TD controls (top panel) and individuals with ASD (bottom panel) in the zero (black), medium (red), and high entropy (blue) conditions at electrode Fz. Stimulus onset is marked by the dotted line.

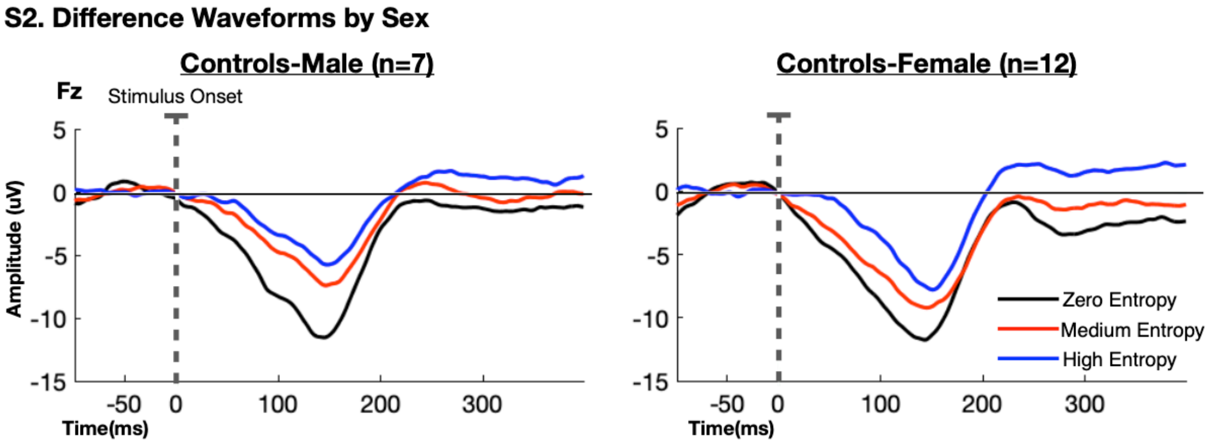


**S2.** Grand-average difference waveforms (deviant tones-standard tones) obtained in male controls (n=7) and female controls (n=12) in the zero (black), medium (red), and high entropy (blue) conditions at electrode Fz. Stimulus onset is marked by the dotted line.

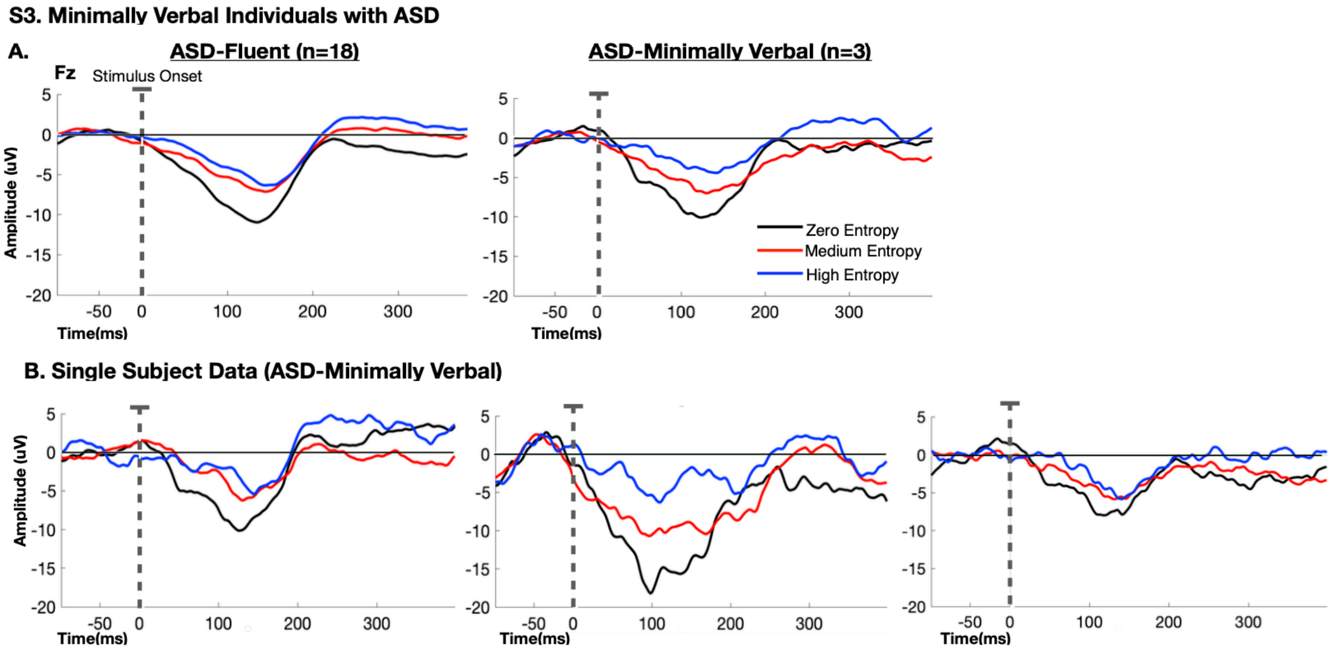


**S3. A)** Grand-average difference waveforms (deviant tones-standard tones) for individuals with ASD who are fluent-speaking (n=18) vs. those who are minimally verbal (n=3) in the zero (black), medium (red), and high entropy (blue) conditions at electrode Fz. **B)** Single subject difference waveforms from the three individuals with ASD who were minimally verbal. Stimulus onset is marked by the dotted line.

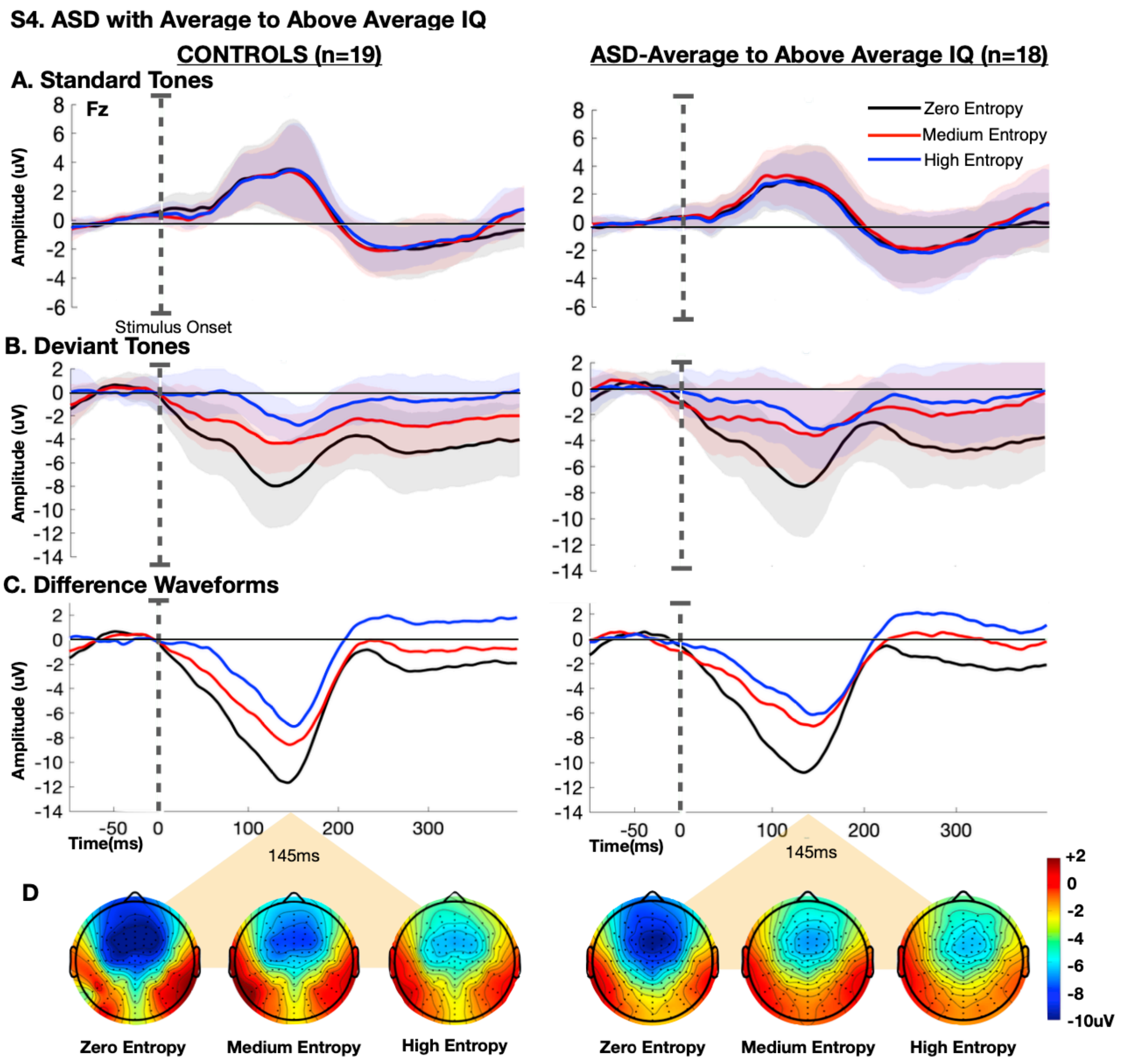


**S4.** Grand-average event related potentials (ERPs) at electrode Fz obtained in neurotypical controls (n=19) and individuals with ASD who have average to above average IQ (n=18) for **A)** standard tones, **B)** deviant tones, and **C)** difference waveforms (deviant tones-standard tones) in the zero (black), medium (red), and high entropy (blue) conditions. Stimulus onset is marked by the dotted line. Shaded regions mark ± 1 SEM. **D)** Topographic representation of the difference between deviant and standard tones at t=145ms for each of the three conditions in controls (left) and individuals with ASD who have average to above average IQ (right).
